# The ‘Stem’ and the ‘Workers’ of the mtDNA population of the cell. Evidence from mutational analysis

**DOI:** 10.1101/2023.04.14.536897

**Authors:** Auden Cote-L’Heureu, Melissa Franco, Yogesh N. K. Maithania, Konstantin Popadin, Dori Woods, Jonathan Tilly, Konstantin Khrapko

## Abstract

Every cell in our body contains a vibrant population of mitochondria, or, more precisely, of mitochondrial DNA molecules (mtDNAs). Just like members of any population mtDNAs multiply (by replication) and ‘die’ (i.e., are removed, either by degradation or by distribution into the sister cell in mitosis). An intriguing question is whether all mitochondria in this population are equal, especially whether some are responsible primarily for reproduction and some - for empowering the various jobs of the mitochondrion, oxidative phosphorylation in the first place. Importantly, because mtDNA is highly damaged such a separation of responsibilities could help greatly reduce the conversion of DNA damage into real inheritable mutations. An unexpected twist in the resolution of this problem has been brought about by a recent high-precision analysis of mtDNA mutations (Sanchez-Contreras et al. 2023). They discovered that certain transversion mutations, unlike more common transitions, are not accumulating with age in mice. We argue that this observation requires the existence of a permanent replicating subpopulation/lineage of mtDNA molecules, which are protected from DNA damage, a.k.a. the ‘stem’ mtDNA. This also implies the existence of its antipode i.e., the ‘worker’ mtDNA, which empowers OSPHOS, sustains damage and rarely replicates. The analysis of long HiFi reads of mtDNA performed by PacBio closed circular sequencing confirms this assertion.

## Introduction: mtDNA damage, mutations, and ‘duplex sequencing’

mtDNA bears high levels of chemical damage due to the genotoxic environment in the mitochondria, which results primarily from byproducts of oxidative phosphorylation, including, but not limited to, reactive oxygen species, (ROS). The effect is apparently exaggerated by poor protection of mtDNA compared to nuclear DNA (lack of nucleosomes) and weaker repair capabilities. This mtDNA damage is converted into mtDNA mutations during mtDNA replication (Marcelino et al. 1998), (Zheng et al. 2006). mtDNA is continuously replicated in the cell: either to compensate for cell proliferation in dividing cells, and to counterbalance mtDNA degradation due to the recycling of mitochondria by mitophagy, which happens even in non-dividing cells. This poses a conceptual problem: if mtDNA is highly damaged and damage is being converted into mutations during replication, now does mtDNA escape a very high mutational rate, expected from converting acute damage into mutations both in somatic cells and even more so in the germline. One possible solution to this enigma is that mtDNA molecules that are being replicated are not damaged, and therefore must be protected from ROS (‘stem’ mtDNA). This logically implies that these mtDNA molecules should probably be located in mitochondria that are less actively involved in oxidative phosphorylation (OXPHOS). Conversely, mtDNA molecules (the ‘workers’) that are actively involved in OXPHOS and therefore highly damaged, should be excluded from replication. There is plenty of evidence of heterogeneity among mitochondria in the cell (Dori), (Döhla et al. 2022),. However, there has been no evidence yet that these heterogeneous subpopulations of mitochondria are *lineal*, i.e. stem mtDNA descends only from stem mtDNA, which is essential for keeping a pure, mutation-free lineage. We have previously reported that mutational patterns in mtDNA mutator PolG mouse imply the possibility of lineal separation of powers between cellular mitochondria (Annis et al. 2018), (Safdar et al. 2016), though other interpretations could not be ruled out.

An unexpected twist to this problem has been brought about by a recent study of mtDNA mutations (Sanchez-Contreras 2023). Traditionally, measuring mtDNA mutations has been challenging. During PCR, damaged nucleotides (chemically altered bases) are replicated to produce mutated daughter molecules (just like they are in the cell). Fortunately, damage-derived mutations are different from real mutations in that they originate from one strand of the parent DNA duplex. The ‘*Duplex sequencing*’, a mutation analysis method, counts only the real mutations by checking whether they are present on both strands of the DNA duplex *before PCR*. (*Sanchez-Contreras 2023)* use duplex sequencing to provide a broad survey of mtDNA mutations in various mouse somatic tissues of different ages. They reported that different types affected the intracellular behavior and fate of a mutation. Note that, paradoxically, this is not the functional type, such as synonymity, but the ‘chemical’ type of the nucleotide change, such as transition vs transversion. Because the ‘chemical’ type of mutation is related to the mechanism of mutagenesis, we reasoned that this result might mean that the fate of mutation (and thus of mtDNA molecule carrying that mutation) depends on the chemical environment in which the molecule is likely to reside, just as predicted by the stem/worker mtDNA hypothesis. To further clarify this relationship, we re-analyzed mutational data reported by (*Sanchez-Contreras 2023)*.

## Methods

The sequencing data used for analysis in this work is publicly available under the accession ID HG002. These reads were generated on the Pacific Biosciences Circular Consensus Sequencing platform and are available as both subreads or HiFi reads, which are preprocessed according to a PacBio standard protocol. The subreads were grouped into consensus reads using the PacBio’s SMRT command line tool ccs. Both types of reads were mapped to a reference genome (human mitochondrion reference genome NC_012920) with another PacBio SMRT tool called pbmm2, which is essentially a wrapper for minimap2. The reads were then aligned to the reference using samtools commands. Finally, variants were called using a custom Python script that returns read ID information, variant position, reference and variant nucleotides, mutant fraction, and quality scores.

The mutational data was analyzed first by counting the number of mutations per molecule, made possible by a simple counting function. Then, a kernel density plot of the number of mutations per molecule was created on the website <u>Wessa.net</u> for visualization. Next, a standard annotation table was used to assign synonymity and mutation type (transition/transversion) to each mutation. By plotting each mutation by type and synonymity, the presence of selective shifts could be compared between each type. Finally, the distribution of quality score for reads that generated mutations was checked and two main groups (“good”/”bad”) were identified based on the average Phred quality score. Then, each quality group was compared by the mutation types that they generated to identify if there was some sort of skew in either direction. All of these analyses were conducted in Excel.

## Results and discussion

### Re-analysis of duplex mutational data leads to the ‘stem/worker’ model of mtDNA population

A novel and exciting discovery of (Sanchez-Contreras 2023) is the high proportion of transversion mutations (C>A/G>T and C>G/G>C) in somatic tissues (compared to many of the previous estimates). Unlike more common transitions, transversions did not accumulate with age and did not clonally expand. These observations are independently broadly confirmed by a more recent, larger duplex sequencing study (Serrano et al. 2023) are rather baffling because mtDNA mutations have an inherent propensity to accumulate with time and to clonally expand within their intracellular populations in a process analogous to the famous genetic drift (Coller, Bodyak, and Khrapko 2002). Of course, expansions of some ‘lucky’ (random) mutations are counterbalanced by convergence and disappearance of a majority of ‘unlucky’ mutations so that the overall mutant fraction remains unchanged as long as the process is truly non-selective. In some mutations, clonal expansion is modulated by selection, positive or negative. Clonal expansion of mtDNA is a fundamentally important process. On the cellular level, expansions allow nascent mutations to reach substantial intracellular mutant fractions and eventually exceed the ‘phenotypic threshold’, which results in the expression of the mutant’s phenotype. On the organismal level, expansion allows nascent germline mtDNA mutations to take over the mtDNA in the germline cell lineage and thus potentially become stably inherited and eventually get fixed in the lineage, the population, and the species.

To make this even more intriguing, another study (Arbeithuber et al. 2020), also using duplex sequencing, discovered high proportion transversions in oocytes. Because mtDNA transversions are known to be inherited only very rarely, high levels of transversions in oocytes imply that transversions must also be somehow excluded from being inherited, probably by the lack of clonal expansion. Thus, the perplexing inability of transversions to expand, noted by (Sanchez-Contreras 2023) in somatic cells, may be an essential property pertinent to mtDNA inheritance.

Why are transversions not subject to otherwise universal clonal expansion and do not accumulate with age as most mtDNA mutations? One obvious possibility is that these mutations are subject to negative selection. However, negative selection must leave a signature – a positive shift in the synonymity of mutations. No such shift was detectable in transversions by (Sanchez-Contreras et al. 2023). We confirmed the lack of synonymity shift by constructing and comparing mutational spectra of synonymous and nonsynonymous mutations (**Figure 1)**.

**Figure 1.**
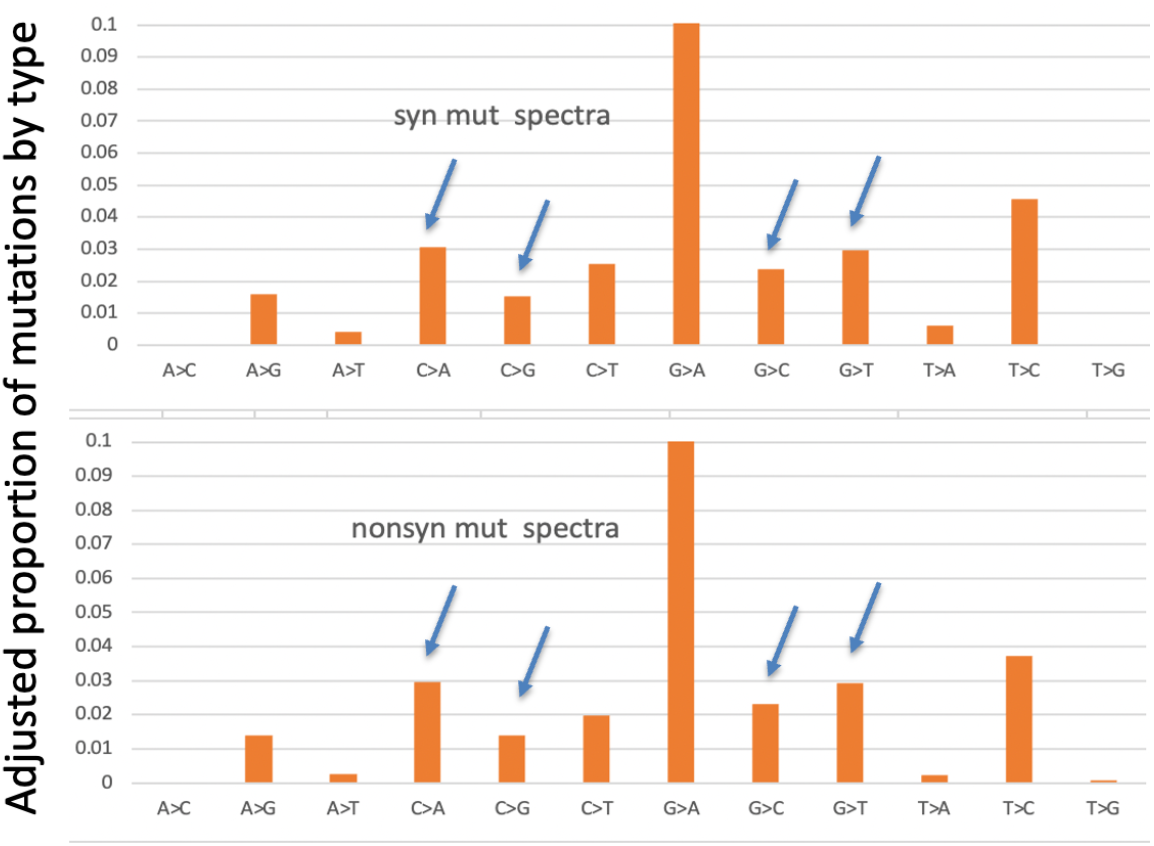
The fraction of transversions (blue arrows) is unchanged between synonymous (top) and non-synonymous (bottom) mutations. Mutational spectra represent the adjusted (divided by the number of available mutable positions) proportion of the mutations by type.

Another way to test for selection are cumulative plots proposed by (Yuan et al. 2020). The results of cumulative curve analysis is shown in **Figure 2**. As one can see, the curves are rather different between G>A (as a representative transition) and C>A (representative transversion), which reflects a much lower propensity of transversions to expand. What is important for our discussion in this context is that the difference (shift) between the synonymous (green) and nonsynonymous (red) mutations is minimal, and even more importantly, the shifts are essentially identical between transitions and transversions. This means that even if there are some selective pressures, they are irrelevant to the transition/transversion disparity. In fact, the direction of the shift (i.e., the red nonsynonymous curve is above the green synonymous curve) implies not negative, but positive selection, or, more likely, a systematically and expectedly higher mutation rates of non-synonymous mutations.

**Figure 2.**
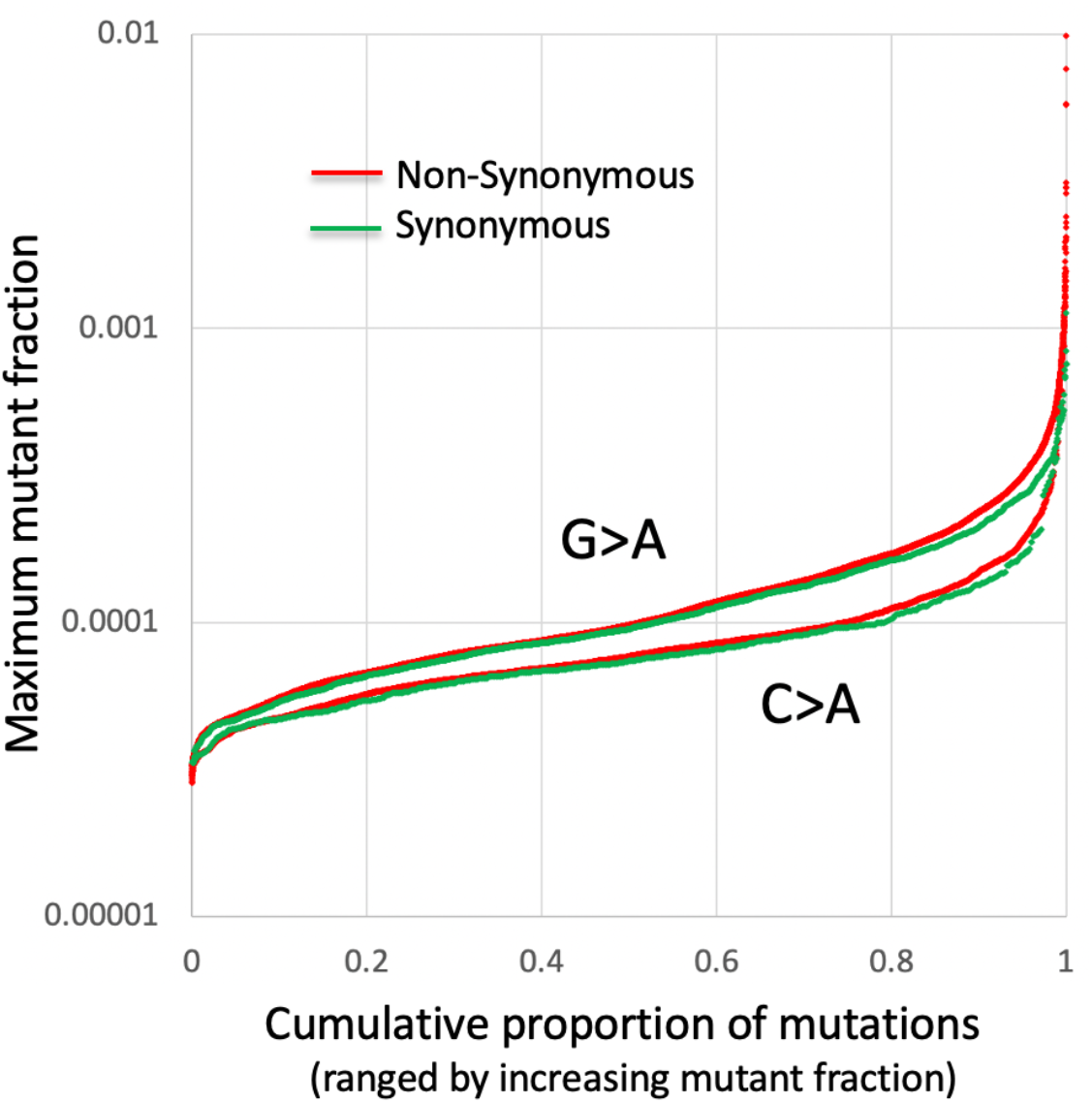
The cumulative proportion graph shows the proportion of mutations that have mutant fraction lower than a certain maximum value. A faster-ascending curve of mutations of a certain category indicates that the mutant fraction of mutations of that category increases more readily, which implies either faster expansion (and therefore selection) or a systematically faster mutation rate.

If transversions are not under direct selection, what else can prevent them from expansion/accumulation? It’s important to note that this cannot be preferential repair proposed by (Sanchez-Contreras 2023). The use of duplex sequencing, implies that these are legitimate double-stranded mutations that therefore cannot be recognized by the DNA repair machinery.

The remaining possibility is that selection is indirect, i.e. whereas transversions are not selected themselves based on their synonymity, but they are preferentially located on those molecules whose replication is impeded or which are destined for removal. This arrangement explains why there is no changes in synonymity. This hypothesis, however, firmly requires that there exists a subpopulation of mtDNA that do not contribute significantly to the genetic pool of the mtDNA population of their cell of residence. Moreover, this subpopulation of molecules should be preferentially targeted by transversion mutations, which are believed to be caused by ROS-induced DNA damage. It is therefore tempting to postulate that this subpopulation is preferentially damaged by ROS products of OXPHS and therefore is located in actively respiring mitochindria. Obviously, these mtDNA fit well the profile of ‘worker’ mtDNA.

To sustain mtDNA population of the cell., there also must exist a subpopulation of molecules that is capable of unlimited replication. These molecules must be relatively depleted of transversions because if transversions were present, they would expand and accumulate, which is not observed. The absence of transversions further implies that the indefinitely replicating subpopulation must be protected from transversion-causing, OXPHOS-related ROS damage DNA damage. Thus these mtDNA must be located in mitochondria with low respiration, which closely follows the profile of ‘stem’ mtDNA.

In conclusion, the logic of our analysis of duplex sequencing mutational database (Sanchez-Contreras et al. 2023), especially the report of non-accumulating transversions inevitably brings us to the ‘stem/worker ‘model of the mtDNA population of the cell.

### HiFi Long read sequencing data from PacBio corroborates the ‘Stem/worker’ model of mtDNA population of the cell

We next set out to test the prediction that there exist more and less damaged mtDNA molecules. Importantly, this issue principally cannot be addressed by duplex sequencing, because the short length of the Illumina reads used in duplex sequencing prevents the analysis of any long-range associations in mtDNA molecules. The latter require monitoring damage along the entire individual mtDNA molecules. This possibility can be realized, for example, by the Pacific Biosciences circular consensus sequencing (CCS) approach in which the entire single mtDNA molecule is repeatedly interrogated by the DNA polymerase molecule performing sequencing by replication, the two strands independently. In this process, DNA damage has been demonstrated to induce sequencing errors specific to the type of damage, although direct relationship between the type of damage and the pattern of errors has not been established (Matsuda, Matsuda, and Yamada 2015). We, therefore, postulated that the frequency of various sequencing errors in PacBio CCS serve as a non-specific proxy of the level of DNA damage on individual DNA molecules.

To this end, we obtained publicly available CCS data, extracted the mtDNA reads and called mutations without the restriction of sequencing quality. Our analysis indeed shows that mtDNA reads fall into a broad range of the number of errors, from no errors to a few dozen, distributed highly non-randomly (**Figure 3**). Additionally, our preliminary observations indicate that a majority of these errors are strand-asymmetric, i.e., originate from one but not the other strand of the DNA duplex. This latter observation corroborates our postulate that sequencing errors represent DNA damage and additionally excludes the possibility that these SNVs originate from NUMTs contaminating mtDNA reads.

**Figure 3.**
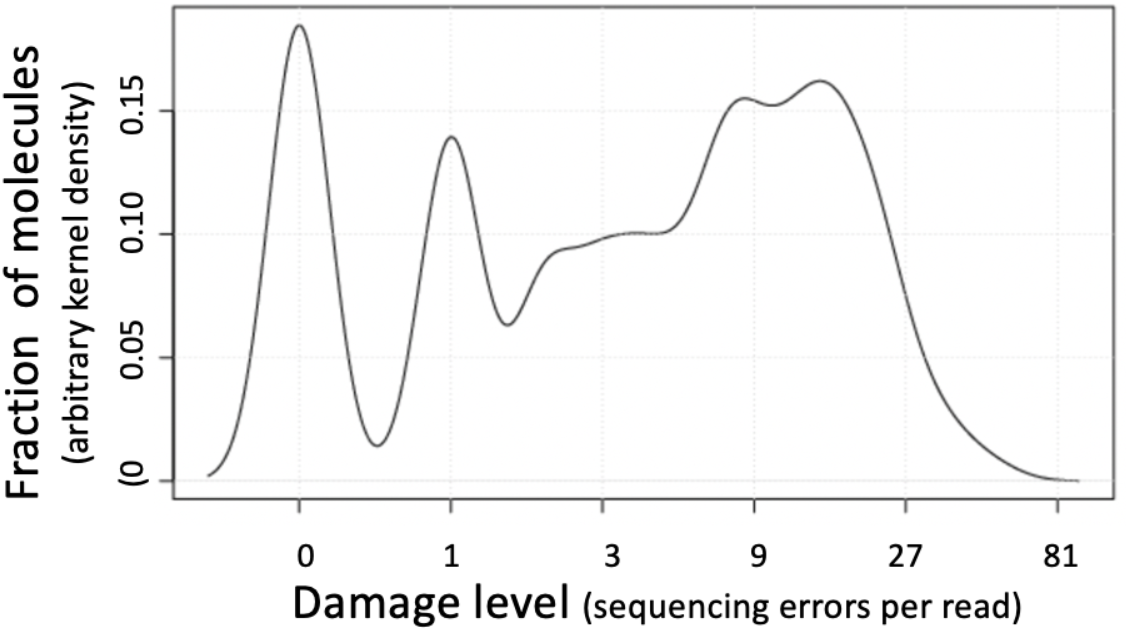
The distribution of DNA damage levels among mtDNA molecules as represented by the number of sequencing errors per HiFi PacBio read. E.g., the sharp peak on the left includes molecules with no errors.

## Discussion

To summarize the results, our analysis consisted of two lines of evidence. First, we looked for an explanation of the puzzling pattern in the recently published database of somatic mtDNA mutations. We showed that the difference in the age-related accumulation of the two types of mutations – transitions (accumulating) and transversions (not accumulation) could only be rationalized in terms of the ‘Stem/Worker’ model (**Figure 4**). We then turned to public data and were able to confirm using lond high-fidelity sequencing reads, that mtDNA molecules indeed consis of subpopulations with low and high levels of damage, which strongly corroborates the model.

**Figure 4.**
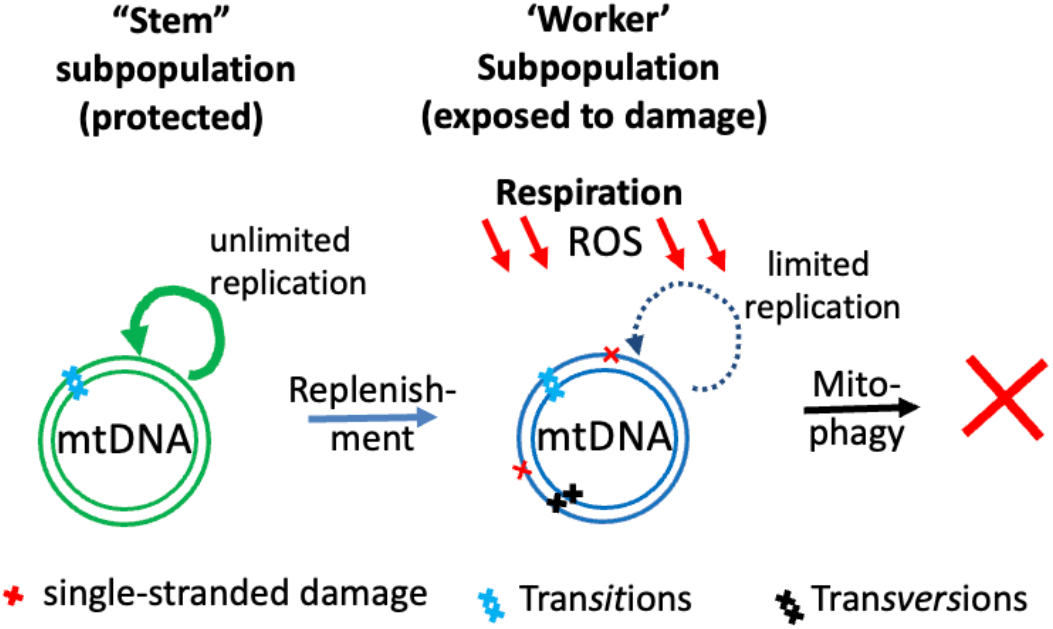
The schematics of the proposed ‘Stem/Worker’ model of the intracellular mtDNA population.

We note that the ‘Stem/Worker’ model in fact is quite probable. In fact, the only essential condition on the model is that there is one way flow: stem mtDNA periodically become on-stem, but non-stem never (very rarely) go back to stem. Stem are protected (e.g., by residing in protected area of the cell). This can be naturally regulated by switching from ‘replicative state’ (protective) to ‘working state” (damaging). Also if something goes bad, like ROS increases, the mitochondrion may cease being stem, and others from stem population replace it). Also, because stem mtDNA are actively replicating, they may be subject to better replication-related repair. This looks like a natural and easy-to-implement scenario. But irrespectively of the mechanism, stem/worker mitochondria model appears to be the only realistic explanation of the transversion behavior in the Sanchez-Contreras et al. manuscript.

The concept of protected stem mitochondria (and it’s opposite - ‘bad’, non-stem, vulnerable subpopulation) provides a reasonable explanation for the reduction of the levels of transversions by ameliorating mitochondrial drugs, as reported by (Sanchez-Contreras et al. 2023). mtDNA molecules that are not stem are subject to damage from which stem mtDNA are protected. ‘Bad’ mtDNAs for that reason *occasionally* carry transversions (or transversion premutagenic lesions). THe damage level in non-stem mtDNAs is naturally at *equilibrium*: non-stem mtDNA get transversion-prone damage (it is partially converted into transversions, and then are eventually removed. In this scenario if one reduces rate of damage (e.g. by the drug treatment), then the load of mutations is proportionally reduced, because the rate of removal remains the same, while the rate of damage is decreased. Equilibrium is shifted. Alternatively (or additionally) these drugs may be ameliorating mitochondrial function by generally increasing turnover rate of mitochondria (reasonable hypothesis), then again it perfectly explains the drop-off of transversions - damage rate is the same, turnover accelerated - equilibrium is shifted in the expected direction.

There are caveats and unresolved issues pertaining to the model, and more research is needed to fully confirm the concept and discern the fine details. For one, transversions mutations could have been in different cells, i.g., in the dying cells. This works fine with kidney and liver and even muscle, but not the heart with eternal cardiomyocytes. But even in heart we can imagine rare ‘bad’ cardiomyocytes which are full of transversions and are eventually eliminated. Best way to test this is to look into individual cells. We do not have single cell analysis for any of somatic tissues. Fortunately, this has been done by the Makova laboratory in oocytes (Arbeithuber et al. 2020). In this study, individual cells appear to contain average numbers of transversions (which argues against the ‘bad cell’ hypothesis. The above logic Ideally needs cell-by-cell analysis and one needs to show that the cell-to-cell variance of transversions is not larger than that of transitions, with the correction to the number of mutants per cell. This will prove that there are no bad cells that are full of transversions (e.g. because they and under constant ROS stress) are destined for elimination.

In conclusion, In this study, we evaluated evidence of the presence, in the population of mtDNA of a cell of two subpopulations, the protected one, broadly responsible for mtDNA replication (‘stem’), and the one responsible for mitochondrial respiration (‘workers’) and therefore exposed to ROS. This separation of function, is potentially of great importance, as it explains how mtDNA, despite high overall damage manages to prevent this damage from being converted into mutations.

